# scRNA + TCR + BCR-seq Reveals the proportion and characteristics of dual TCR T cells and dual BCR B cells in mice infected with Echinococcus granulosus

**DOI:** 10.1101/2025.07.24.666567

**Authors:** Kai Quan, Peng Su, Hongxia Yang, Yadong Gong, Xinsheng Yao

**Author notes:** Co-first Author (Kai Quan, Peng Su).

## Abstract

Cystic echinococcosis (CE) is a zoonotic parasitic disease caused by Echinococcus granulosus, and the T and B cell responses and regulatory mechanisms in CE remain unclear. In this study, we analyzed the data of peripheral blood and spleen of mice infected with Echinococcus granulosus(EG) by scRNA+TCR+BCR-seq, and made the following novel findings: (1) Peripheral blood of Echinococcus granulosus (EG)-infected mice contained a higher proportion of dual TCR(T cell receptor) T cells with higher clonality than uninfected mice; (2) There was a bias in V gene usage between single TCR T and dual TCR T cells, as well as between single BCR(B cell receptor) B cells and dual BCR B cells in EG-infected mice; (3) Dual TCR Tregs, dual TCR effector CD8 T cells, dual BCR Bregs, and dual BCR memory/dividing B cells exhibited relatively high clonal expansion in EG-infected mice; (4) Shared CDR3 sequences and consistently highly expressed mRNA molecules exist between single receptor T/B and dual receptor T/B cells in EG-infected mice. KEGG enrichment analysis revealed that dual BCR B cells are closely related to the expression of factors associated with pathways such as “Th1/Th2 differentiation” and “antigen processing and presentation”. These results suggest that dual TCR T and dual BCR B cells may be involved in the specific responses and regulation of EG-infected mice, providing a novel perspective for the adaptive immune response mechanisms and targeted therapy of parasitic infections.

**Author summary:** Echinococcosis is a zoonotic disease currently prevalent mainly in various parts of Africa. At present, it cannot be completely cured; treatment relies solely on medication to alleviate symptoms. The immunoregulatory mechanisms underlying the disease remain unclear. Using single-cell RNA sequencing, we have identified a novel T- and B-cell subpopulation that may play a role in the immune response to the parasite. This finding opens a new avenue for researchers investigating parasitic diseases.

## 1. Introduction

Cystic echinococcosis (CE) is one of the 17 neglected tropical diseases closely monitored by the World Health Organization, primarily caused by infection with Echinococcus granulosus, affecting over one million people. CE can damage multiple organs in the human body and is typically a lifelong disease that threatens life, making it difficult to prevent, control, and treat, resulting in annual costs exceeding 3 billion US dollars[1, 2]. CE has become a global public health issue. The specific T and B cell immune responses and regulatory mechanisms in CE have not been fully elucidated. Although some progress has been made in vaccine research[3, 4], potentially effective therapeutic drugs are still awaiting clinical trials[5].

During the life cycle of Echinococcus granulosus (EG), it develops into the larval stage within the intermediate host, known as hydatid cysts. However, it does not further develop into adults within the intermediate host. These larval stages of hydatid cysts can produce eggs or larvae that can infect new hosts. The larval stage can induce an immune evasion response in the host, which helps the parasite survive long-term within the host and avoid clearance by the host’s immune system[6, 7]. In the early stages of EG infection in mice, the immune response in vivo is predominantly Th1, but as the disease progresses to the middle and late stages, Th2 cells become dominant[8]. IFN may play a protective role in Echinococcus multilocularis infection[9]. The cytokine profiles favoring Th1 shift and lesions were significantly reduced in patients with hepatitis C and CE treated with IFN-α[10], suggesting that IFN-α contributes to immune protection against echinococcus infection and slows disease progression. Zhang S et al. also found that the combination of albendazole (ABZ) and IFN-α may be effective in treating CE in humans and animals[11]. Treg cells proliferate during EG infection, and targeting Foxp3+ Treg cells may be effective[12]. Breg cells proliferate in the late stages of EG infection and may regulate specific immune responses by modulating cytokine expression and inducing the secretion of inhibitory cytokines IL-10 and TGF-β[13].

The clonal selection theory, which posits that “a lymphocyte expresses only one type of antigen receptor,” has been challenged and supplemented by the existence of “dual TCR T and dual BCR B cells”[14–16]. Since the first detection of two functional Vα and Vβ gene rearrangements in a single T cell at the mRNA level in 1988[17, 18], multiple laboratories have identified the presence of dual TCR T and dual BCR B cells[19]. In recent years, it has been confirmed that dual TCR T and dual BCR B cells are closely related to the occurrence and development of tumors, infections, and autoimmune diseases[20–27].

The antigenic epitopes and virulence factors of parasites differ from those of bacteria and viruses. Research on dual-receptor lymphocytes following parasitic infections has not yet been reported. This study innovatively found a higher proportion and clonal expansion of dual TCR T and dual BCR B cells in EG-infected mice through detailed comparative analysis of scRNA + TCR + BCR-seq shared data from EG-infected mice. The CDR3(complementarity-determining region 3) composition, source subgroups, and highly expressed mRNA molecules of these cells suggest that they may be involved in the specific immune responses and regulation of EG infection, providing a novel perspective for the adaptive immune response mechanisms and targeted therapy of parasitic infections.

## 2. Materials and Methods

### 2.1 Study Subjects and Samples

The study subjects and samples in this study were derived from the article published by Wu J et al. ( Single-Cell·RNA Sequencing Reveals Unique Alterations in the Immune Panorama and Treg Subpopulations in Mice during the Late Stages of Echinococcus granulosus Infection , Infection and Immunity,Epub 2023 Apr 11,doi:10.1128/iai.00029-23). The analyzed data were obtained from the scRNA + TCR + BCR-seq sequences shared by the authors (Gene Expression Omnibus database, accession numbers: GSE216347 and GSE225312, https://www.ncbi.nlm.nih.gov/geo/query/acc.cgi?acc=GSE216347, https://www.ncbi.nlm.nih.gov/geo/query/acc.cgi?acc=GSE225312).

#### Study subjects

Wild type 6 to 8-week-old female BALB/c mice were used (6 mice in total), divided into EG-infected group (3 mice) and non-infected group (3 mice). Hydatid cysts isolated from patients with confirmed CE were injected intraperitoneally into the mice. The infected mice were euthanized at the late stage of infection (week 22). The non-infected group was injected with PBS as a control.

#### Samples

Peripheral blood and spleens were collected from both EG-infected and non-infected mice for scRNA + TCR + BCR-seq sequencing. The samples were divided into four groups: IP = peripheral blood samples from EG-infected mice; UP = peripheral blood samples from non-infected mice; IS = spleen samples from EG-infected mice; US = spleen samples from non-infected mice. Sample names, accession numbers, total number of T and B cells sequenced, and total number of TCR and BCR pairs are shown in Figures 1-C and 1-D.

**Figure 1.**
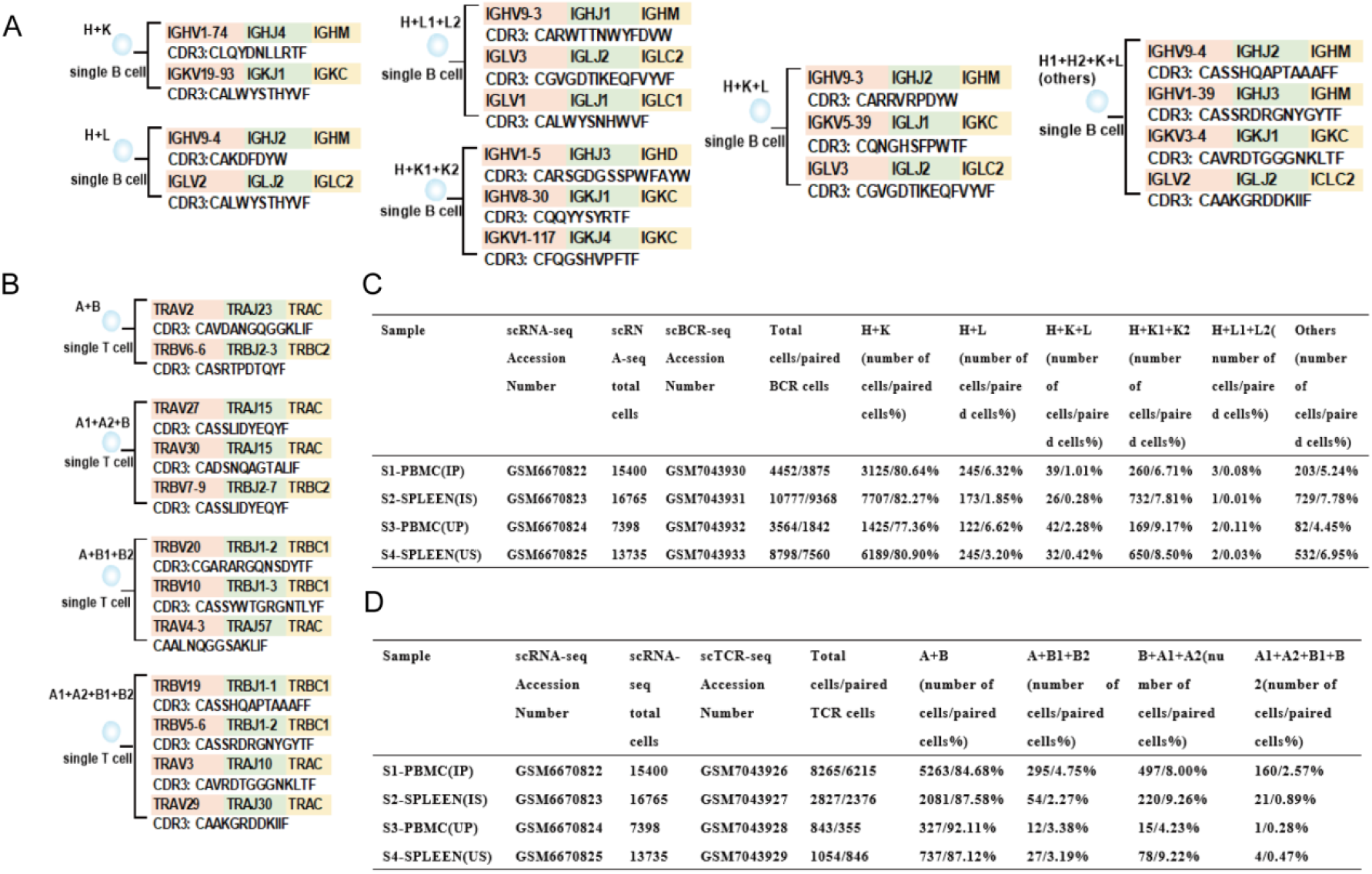
Examples of CDR3 sequences from single and dual TCR T cells and single and dual BCR B cells, along with sample names, shared data serial numbers, and counts and proportions of TCR and BCR pairing types. A. Examples of CDR3 sequences for different pairing types of BCR heavy (H) chains, kappa (K) chains, and lambda (L) chains (H+K; H+L; H+K1+K2; H+K+L; H+L1+L2; others). B. Examples of CDR3 sequences for different pairing types of TCR alpha (α) chains and beta (β) chains (A+B; A1+A2+B; A+B1+B2; A1+A2+B1+B2). C. Sample names; shared serial numbers for single-cell RNA sequencing (scRNA-seq); total count of functional B cells; shared serial numbers for single-cell BCR sequencing (scBCR-seq); total count of pairable BCR B cells; counts and proportions of cells with different BCR pairing types. D. Sample names; shared serial numbers for single-cell RNA sequencing (scRNA-seq); total count of functional T cells; shared serial numbers for single-cell TCR sequencing (scTCR-seq); total count of pairable TCR T cells; counts and proportions of cells with different TCR pairing types.

### 2.2 Analysis Workflow for Single BCR B Cells and Dual BCR B Cells

scBCR-seq data analysis workflow: (1) Sequences marked as “FALSE” in the “is_cell” and “high_confidence” columns were deleted; sequences in the “Chain” column that were not “IGH,” “IGK,” or “IGL” were excluded; sequences marked as “Non” or “FALSE” in the “Productive” column were deleted; B cells with only one functional chain sequenced were removed. (2) In a single B cell, two functional chains can be paired and assembled into a single BCR B cell, referred to as BCR Heavy chain (H) + BCR light chain (Kappa or Lambda). In a single B cell, three functional chains can be paired and assembled into a dual BCR B cell, referred to as H + K1 + K2; H + L1 + L2; H + K + L. In a single B cell, four (or more) functional chains can be paired and assembled into a BCR B cell, referred to as others. Two or more BCR pairing types are collectively named dual BCR B cells. For each BCR pairing type, one B cell was selected, and the organization of the V(D)J gene family and the CDR3 amino acid sequence were listed (Figure 1-A).

### 2.3 Analysis Workflow for Single TCR T Cells and Dual TCR T Cells

scTCR-seq data analysis workflow: (1) Sequences marked as “FALSE” in the “is_cell” and “high_confidence” columns were deleted; sequences in the “Chain” column that were not “TRA” or “TRB” were excluded; sequences marked as “Non” or “FALSE” in the “Productive” column were deleted; T cells with only one functional chain sequenced were removed. (2) In a single T cell, two functional chains can be paired and assembled into a single TCR T cell, referred to as TCR alpha chain (A) + TCR beta chain (B). In a single T cell, three functional chains can be paired and assembled into a dual TCR T cell, referred to as A + B1 + B2; B + A1 + A2. In a single T cell, four functional chains can be paired and assembled into a multi-TCR T cell, referred to as A1 + A2 + B1 + B2. Two or more TCR pairing types are collectively named dual TCR T cells. For each TCR pairing type, one T cell was selected, and the organization of the V(D)J gene family and the CDR3 amino acid sequence were listed (Figure 1-B).

### 2.4 CDR3 Repertoire Characterization of scBCR-seq and scTCR-seq

After quality control of scTCR + BCR-seq data, the R package immunarch was used to compare and analyze the CDR3 repertoire characteristics of single TCR T, dual TCR T, single BCR B, and dual BCR B cells: (1) Clonality: The proportion of cells with clonal counts ≥2 and clonal counts =1 were compared among single TCR T, dual TCR T, single BCR B, and dual BCR B cells; (2) V gene usage bias: The usage of V gene subfamilies was compared among single TCR T, dual TCR T, single BCR B, and dual BCR B cells, with statistical analysis of V gene usage differences using SPSS; (3) Overlap: The overlap of CDR3 sequences was analyzed using the R package immunarch; (4) Each T and B cell’s unique barcode was used to integrate scRNA-seq and scTCR + BCR-seq data for analysis.

### 2.5 scRNA-seq Data Quality Control and Integration with scTCR + BCR-seq

The scRNA-seq data shared in GEO were imported into R for analysis: (1) The harmonyR package was used for sample integration, and cells with more than 12,000 or fewer than 200 unique genes were excluded from the analysis; (2) Cells with mitochondrial gene percentages exceeding 5% and log10GenesPerUMI (genes per UMI) less than 0.8 were removed from the dataset, and only genes expressed in 10 or more cells were used for further analysis; (3) Doublets were identified and removed using the “DoubletFinder” R package in conjunction with the “Seurat” R package version 5; (4) Normalization was performed with the normalization method LogNormalize in the Seurat package (version 4.0.1) and with a scale factor with a default value of 10,000; (5) The Seurat object’s ScaleData, FindVariableFeatures, and RunPCA functions were analyzed, and dimensionality reduction clustering was performed using the RunUMAP function; the top 10 mRNA expressions of each sample subgroup were selected using the FindAllMarkers function; the “SingleR” package was used to annotate cell types, and the clustering results were corrected based on the expression of canonical genes; (6) T cells were clustered based on the expression of canonical markers and cell subgroups: EffectorCD8T (Cd8a, Gzmk, Emoes, Ly6c1, Ly6c2, Nkg7); NaiveCD8T (Cd8a, Sell, Ccr7); NaiveCD4T (Cd4, Sell, Ccr7); IFN_CD4 (Cd4, Isg15); Th1 (Cd4, Tbx21, Stat4, Ifr8); Th2 (Cd4, Stat6, Il-4); Treg (Cd4, Foxp3); (7) B cells were clustered based on the expression of canonical markers and cell subgroups: Naive_B (Cd19, Ighd); Memory_B (Cd19, Ighd, Cd27); Breg (Cd19, Cd1d1, Cd1d2); Plasma (Cd19, Jchain, Xbp1); Dividing_B (Cd19, Mki67, H2afz, Cdk1, Aicda); (8) Data that met both scRNA-seq quality control and scTCR + BCR-seq screening criteria were integrated for analysis based on each cell’s unique barcode.

### 2.6 Major Tools, Software, and Statistical Methods

Software packages: The R packages “Seurat,” “ggplot2,” “DoubletFinder,” Adobe, and GraphPad Prism (version 5) were used for data visualization; data analysis was performed using R studio (v3.3.3) and GraphPad Prism (version 5); the “SingleR” package was used for cell type annotation, with manual correction based on canonical gene expression; Statistical methods: IBM SPSS Statistics 26 software was used for statistical analysis; when n ≥ 40 and Tmin ≥ 5, Pearson’s chi-square test was used to compare multiple ratios between two groups; when n < 40 or Tmin < 1, Fisher’s exact test was used.

## 3. Results

### 3.1 Proportion and Pairing Characteristics of Dual TCR T and Dual BCR B Cells

Dual TCR T and dual BCR B cells were identified in the spleen and peripheral blood of EG-infected mice, with CDR3 sequences shown in Figures 1-A and 1-B. The proportions of dual BCR B cells in the peripheral blood and spleen of EG-infected mice were 13.07% and 15.88%, respectively, with the main pairing type being H + K1 + K2 (6.71% in peripheral blood; 7.81% in spleen) (Figure 1-C). The proportions of dual TCR T cells in the peripheral blood and spleen of EG-infected mice were 15.32% and 12.42%, respectively, with the main pairing type being A1 + A2 + B (8.00% in peripheral blood; 9.26% in spleen) (Figure 1-D).

#### Dual TCR T cells

IP group (15.32%) > UP group (7.89%); no difference between US (12.88%) and IS groups (12.42%) (Figure 2-A); IP group (15.32%) > IS group (12.42%) (Figure 2-C).

**Figure 2.**
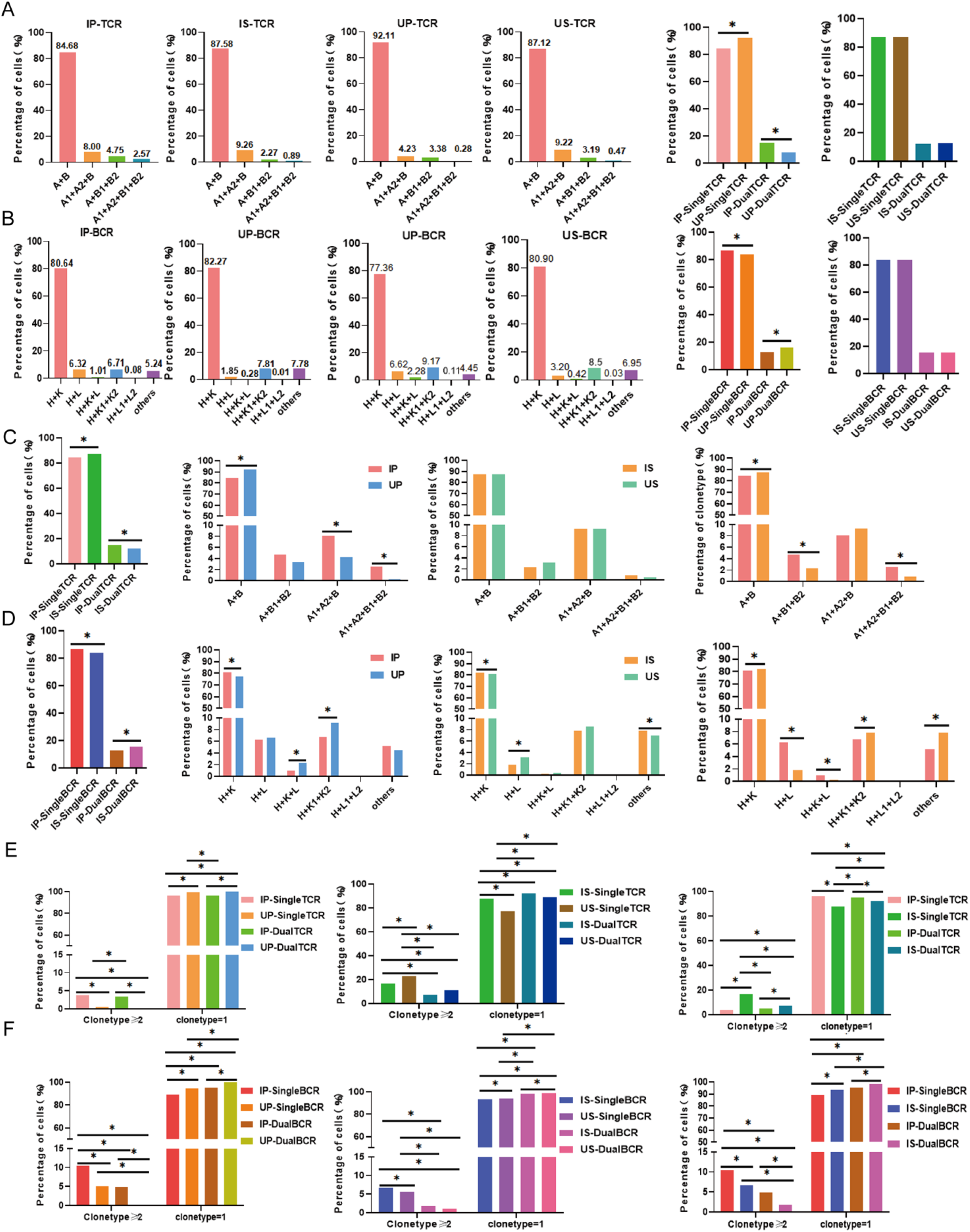
Analysis of TCR and BCR CDR3 pairing types and clonality in IP, IS, UP, and US using scTCR-seq and scBCR-seq. A. Proportions of single TCR T and dual TCR T pairing types in IP, IS, UP, and US, with comparisons between total single TCR T and total dual TCR T (IP vs. UP; IS vs. US). B. Proportions of single BCR B and dual BCR B pairing types in IP, IS, UP, and US, with comparisons between total single BCR B and total dual BCR B (IP vs. UP; IS vs. US). C. Comparison of the proportions of total single TCR T and total dual TCR T cells between the IP and IS groups; comparison of TCR pairing types among IP, IS, UP, and US groups. D. Comparison of the proportions of total single BCR B and total dual BCR B cells between the IP and IS groups; comparison of BCR pairing types among IP, IS, UP, and US groups. E. Comparison of clonal expansion (clonetype ≥ 2) of single TCR T and dual TCR T cells among IP, IS, UP, and US. F. Comparison of clonal expansion (clonetype ≥ 2) of single BCR B and dual BCR B cells among IP, IS, UP, and US.(^*^: P > 0.05)

#### Dual BCR B cells

UP group (16.02%) > IP group (13.03%); no difference between US group (15.90%) and IS group (15.88%) (Figure 2-B); IS group (15.88%) > IP group (13.03%) (Figure 2-D).

Among the four groups, the dual TCR T cell pairing type with the highest proportion was A1 + A2 + B, in the order of IS (9.26%) > US (9.22%) > IP (8.00%) > UP (4.23%). For dual BCR B cells, the most common pairing type was H + K1 + K2, with proportions in the order of UP (9.17%) > US (8.50%) > IS (7.81%) > IP (6.71%) (Figure 2-C and 2-D).

### 3.2 Clonality of CDR3 Repertoire

Clonal expansion (clonetype ≥2) proportions were compared as follows: (1) IP single TCR (3.84%) > IP dual TCR (3.47%) > UP single TCR (0.62%) > UP dual TCR (0%); US single TCR (22.93%) > IS single TCR (16.97%) > US dual TCR (11.01%) > IS dual TCR (7.46%); IS single TCR (16.97%) > IS dual TCR (7.46%) > IP single TCR (3.84%) > IP dual TCR (3.47%) (Figure 2-E). (2) IP single BCR (10.53%) > UP single BCR (5.11%) > IP dual BCR (4.95%) > UP dual BCR (0%); IS single BCR (6.70%) > US single BCR (5.70%) > IS dual BCR (1.75%) > US dual BCR (1.16%); IP single BCR (10.53%) > IS single BCR (6.70%) > IP dual BCR (4.95%) > IS dual BCR (1.75%) (Figure 2-F).

### 3.3 V Gene Usage in CDR3 Repertoire

In the IP group, dual TCR T cells preferentially used TRAV3 and TRBV17, while single TCR T cells preferentially used TRAV5 and TRBV2. Dual BCR B cells preferentially used IGHV8, while single BCR B cells preferentially used IGHV1 (Supplementary Figure 1-A). In the IS group, dual TCR T cells preferentially used TRAV1 and TRBV10, while single TCR T cells preferentially used TRAV11. Single BCR B cells preferentially used IGHV9 (Supplementary Figure 1-B).

Compared to the UP group, dual TCR T cells in the IP group preferentially used TRBV19. Compared to the US group, dual TCR T cells in the IS group preferentially used TRBV12. Dual BCR B cells preferentially used IGHV8 and IGHV14. When comparing the IP and IS groups, dual TCR T cells in the IP group preferentially used TRBV5, while dual TCR T cells in the IS group preferentially used TRAV1 and TRBV2. No significant differences were observed in V gene usage between dual BCR B cells in the IP and IS groups (Supplementary Figures 1-C,1-D,1-E).

### 3.4 Dual TCR T and Dual BCR B Cells in Mouse Peripheral Blood Samples

In EG-infected mice, **T cell subgroups were clustered as follows:** Effector_CD8T; IFN_CD4T; Naïve_CD4T; Naïve_CD8T; Th1; Th2; Treg (Figure 3-A). Dual TCR T cells were found in each subgroup. **The sources of dual TCR T cells were as follows:** Naïve_ (72.77%) > Naïve_CD8T (12.90%) > Treg (5.29%) > IFN_CD4T (4.77%) > Effector_CD8T (3.10%) > Th2 (0.90%) > Th1 (0.26%). Clonal expansion was observed in dual TCR T cells within the Effector_CD8T, Naïve_CD4T, and Naïve_CD8T subgroups, with the highest levels in Effector_CD8T. Similarly, single TCR T cells exhibited the highest clonal expansion in Effector_CD8T (Figure 3-B). B cell subgroups were clustered as follows: Breg; Plasma; Naïve_B; Memory_B (Figure 3-A). Dual BCR B cells were found in each subgroup. The sources of dual BCR B cells were as follows: Naïve_B (57.73%) > Plasma (17.11%) > Memory_B (14.85%) > Breg (10.31%). Clonal expansion was observed in single BCR B cells, with the highest levels in Plasma cells. Dual BCR B cells exhibited clonal expansion in Naïve_B and Memory_B cells, with no significant differences between the two (Figure 3-C).

**Figure 3.**
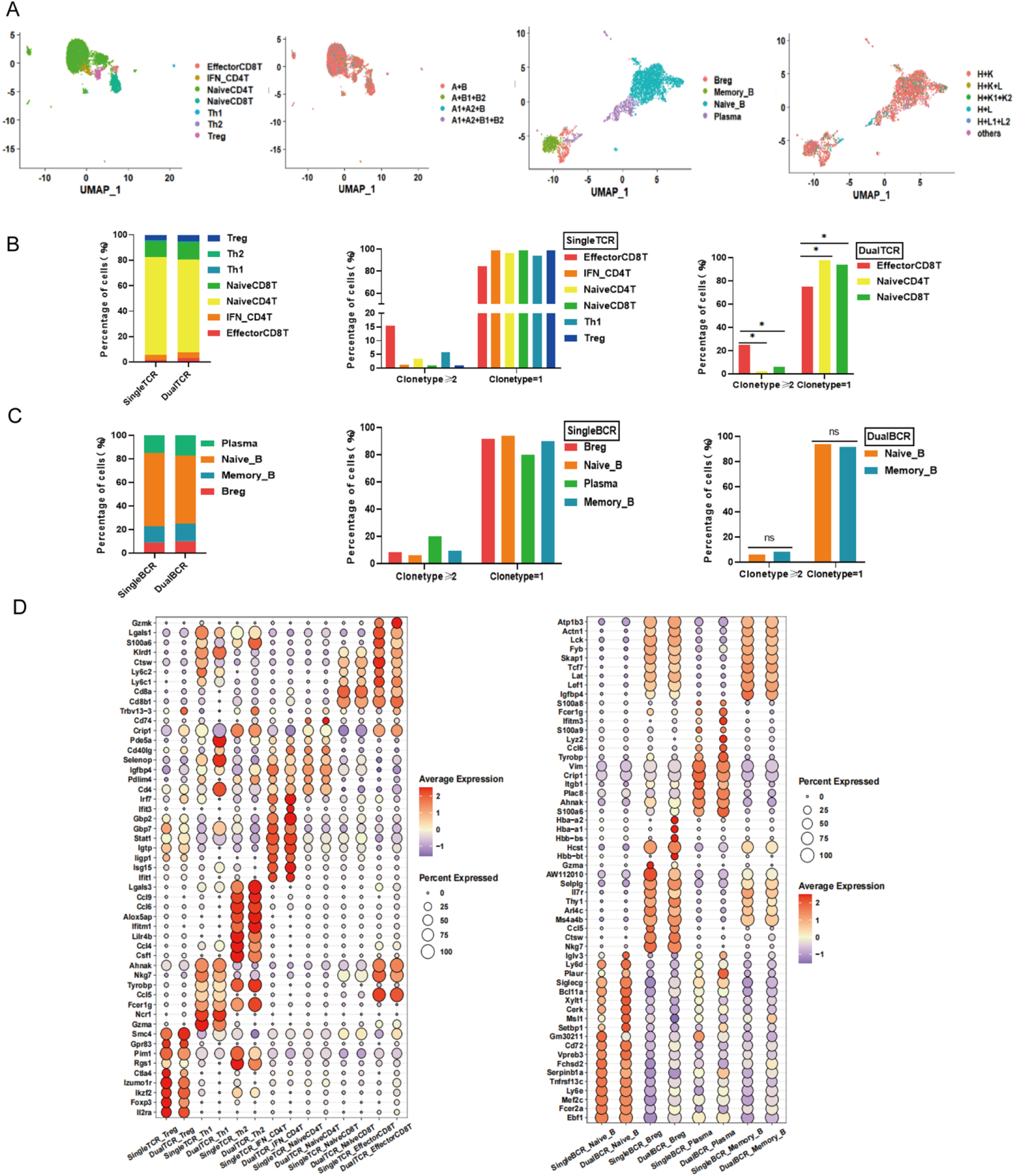
Analysis of T and B cell subsets, clonality, and mRNA expression in the IP group using ScRNA+TCR+BCR-seq. A. UMAP clustering maps of T cell and B cell subsets, with distribution charts of TCR and BCR pairing types. B. Comparison of the distribution and clonal expansion (clone type ≥ 2) of single TCR T cell and dual TCR T cell subsets. C. Comparison of the distribution and clonal expansion (clone type ≥ 2) of single BCR B cell and dual BCR B cell subsets. D. Comparison of the top 10 mRNA expressions in single TCR T and dual TCR T cells, and comparison of the top 10 mRNA expressions in single BCR B and dual BCR B cells. (*: P > 0.05)

Characteristic gene and cytokine expression analysis between single TCR T and dual TCR T cell subgroups: The expression profiles of single TCR T cell and dual TCR T cell subgroups were generally consistent. However, some subgroups exhibited higher expression of characteristic genes. For example, dual TCR Th2 cells had higher expression of the characteristic gene Il-4 compared to single TCR Th2 cells, and dual TCR IFN_CD4T cells had higher expression of Isg15, Isg20, and Ifit3 compared to single IFN_CD4T cells (Supplementary Figure 2-A). Differential expression factors between single BCR B and dual B cell subgroups: The expression profiles of single BCR B cell and dual BCR B cell subgroups were also generally consistent. However, some subgroups exhibited higher expression of characteristic genes. For example, dual BCR Breg cells had higher expression of the characteristic gene Cd1d1 compared to single TCR Th2 cells (Supplementary Figure 2-A).

In uninfected mice, dual TCR T cells were found in the following subgroups: Naïve_CD4T; Naïve_CD8T; Treg; Effector_CD8T. No clonal expansion was observed in dual TCR T cells. Dual BCR B cells were found in the following subgroups: Plasma; Naïve_B; Memory_B. No clonal expansion was observed in dual BCR B cells (Supplementary Figures 4-A,4-B,4-C).

### 3.5 Dual TCR T and Dual BCR B Cells in Spleen Samples

In EG-infected mice, **T cell subgroups were clustered as follows:** Effector_CD8T; IFN_CD4T; Naïve_CD4T; Naïve_CD8T; Th2 (Figure 4-A). Dual TCR T cells were found in each subgroup. **The sources of dual TCR T cells were as follows:** Naïve_CD4T (36.99%) > Treg (21.92%) > Naïve_CD8T (17.35%) > IFN_CD4T (12.79%) > Effector_CD8T (6.85%) > Th2 (4.11%). Clonal expansion was observed in dual TCR T cells within the Effector_CD8T, IFN_CD4T, and Treg subgroups, with the highest levels in Effector_CD8T. Similarly, single TCR T cells exhibited the highest clonal expansion in Effector_CD8T and IFN_CD4T (Figure 4-B). B cell subgroups were clustered as follows: Breg; Plasma; Naïve_B; Memory_B; Dividing_B (Figure 4-A). Dual BCR B cells were found in each subgroup. The sources of dual BCR B cells were as follows: Naïve_B (73.44%) > Breg (13.74%) > Dividing_B (8.26%) > Plasma (3.99%) > Memory_B (0.56%). Clonal expansion was observed in single BCR B cells, with the highest levels in Plasma cells. Dual BCR B cells exhibited clonal expansion primarily in Dividing_B cells (Figure 4-C).

**Figure 4.**
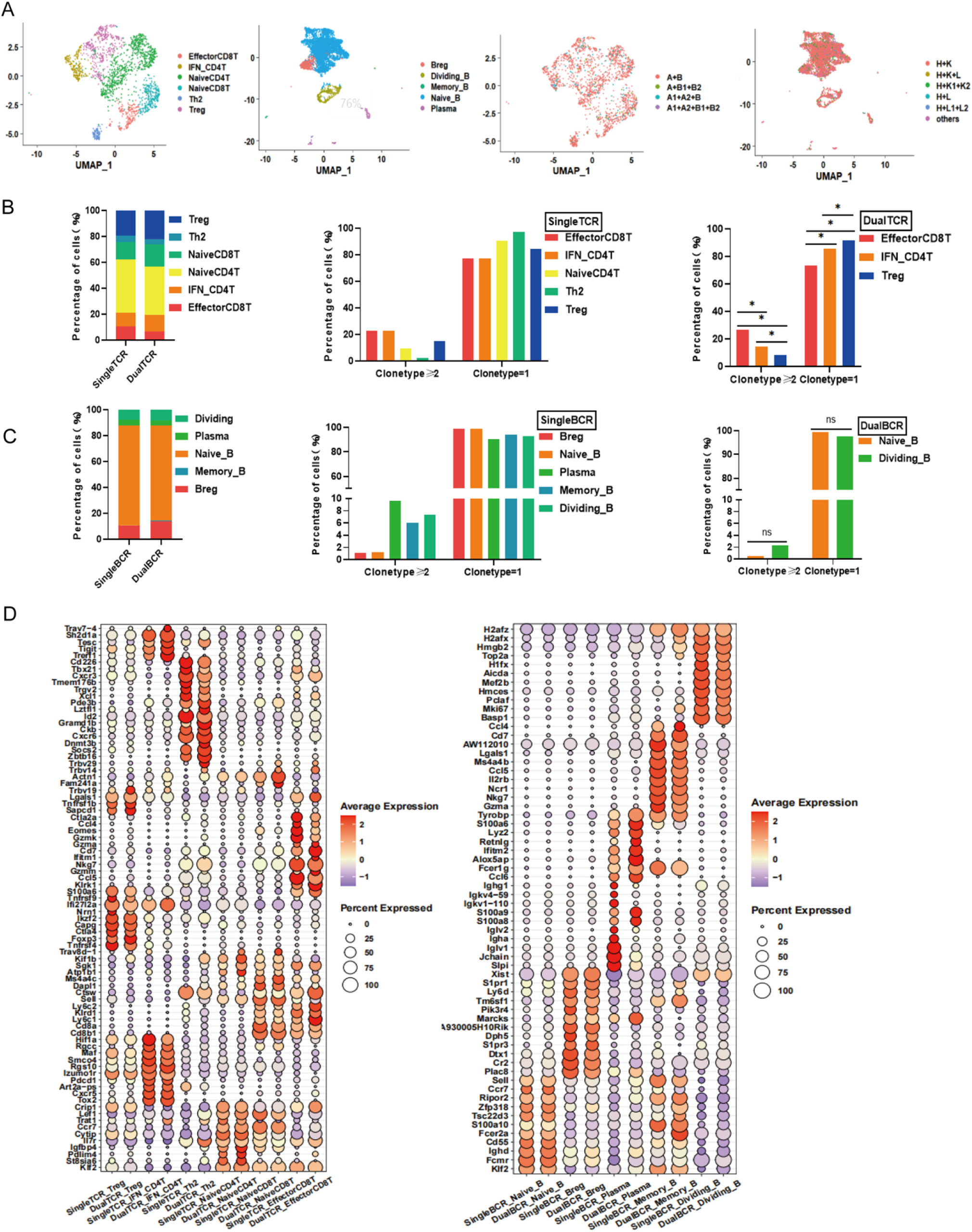
Analysis of T and B cell subsets, clonality, and mRNA expression in the IS group using ScRNA+TCR+BCR-seq. A. UMAP clustering plots of T cell and B cell subsets; distribution plots of TCR and BCR pairing types. B. Comparison of the distribution and clonal expansion (clone type ≥ 2) among single TCR T cell and dual TCR T cell subsets. C. Comparison of the distribution and clonal expansion (clone type ≥ 2) among single BCR B cell and dual BCR B cell subsets. D. Comparison of the top 10 mRNA expressions in single TCR T and dual TCR T cells, and the top 10 mRNA expressions in single BCR B and dual BCR B cells. (^*^: P > 0.05)

Analysis of characteristic gene and cytokine expression between single TCR T and dual TCR T cell subgroups: The expression profiles of single TCR T cells and dual TCR T cell subgroups were generally consistent. However, some subgroups exhibited higher expression of characteristic genes. For example, dual TCR Treg cells had higher expression of the inhibitory gene Foxp3 compared to single TCR Treg cells (Supplementary Figure 3-A). Differential expression factors between single BCR B and dual B cell subgroups: The expression profiles of single BCR B cells and dual BCR B cell subgroups were generally consistent (Supplementary Figure 3-A).

In the IS and US groups, significant differences in expression were observed for PDCD1, Clta4, Lag3, and Havcr2 in the Treg subgroup.

In uninfected mice, **dual TCR T cells were found in the following subgroups:** Naïve_CD4T; Naïve_CD8T; Treg; Effector_CD8T. Clonal expansion was observed primarily in dual TCR Naïve_CD4T cells. Dual BCR B cells were found in the following subgroups: Plasma; Naïve_B; Memory_B. Clonal expansion was observed only in dual BCR Naïve_B cells (Supplementary Figures 5-A,5-B,5-C).

### 3.6 Top 10 mRNAs of Single-Receptor and Dual-Receptor T/B Cells; Shared CDR3 Sequences; Top 10 mRNAs and KEGG Analysis of Total Dual TCR T and Dual BCR B Cells

In EG-infected mice, the top 10 mRNA molecules expressed in each single TCR T and dual TCR T subgroup, as well as each single BCR B and dual BCR B subgroup, were generally consistent (Figures 3-D, 4-D). In uninfected mice, the consistency of the top 10 mRNA molecules expressed in each single TCR T and dual TCR T subgroup, as well as each single BCR B and dual BCR B subgroup, was lower than that observed in EG-infected mice (Supplementary Figures 4-D, 5-D).

In both EG-infected and uninfected mice, partial sharing of CDR3 sequences was observed between single TCR T cell subgroups, dual TCR T cell subgroups, and between single and dual T cell subgroups. The number of shared CDR3 sequences was higher in EG-infected mice compared to uninfected mice. Similarly, partial sharing of CDR3 sequences was observed between single BCR B cell subgroups, dual BCR B cell subgroups, and between single and dual B cell subgroups, with higher numbers of shared CDR3 sequences in EG-infected mice (Supplementary Figures 2-B, 3-B, 4-E, 5-E).

Differences in the expression of the top 10 mRNA molecules were observed between the spleen and peripheral blood in both EG-infected and uninfected mice for both total dual TCR T and total dual BCR B cells (Supplementary Figure 6). KEGG enrichment analysis of total dual TCR T and total dual BCR B cells in EG-infected mice revealed that both single and dual BCR B cells in peripheral blood significantly expressed genes related to “Th1 and Th2 differentiation” and “Antigen processing and presentation.” In the spleen, single and dual BCR B cells significantly expressed genes related to “Intestinal immune network for IgA production” and “Antigen processing and presentation” (Supplementary Figures 7, 8).

## 4. Discussion

The specificity, diversity, and memory of T and B cell responses to antigens are based on the clonal selection theory, which posits that “a lymphocyte expresses only one type of antigen receptor.”[28]However, incomplete V(D)J allelic exclusion recombination can lead to the existence of “dual receptor lymphocytes.” In recent years, dual TCR T and dual BCR B cells have been shown to be closely related to the occurrence and development of tumors, infections, and autoimmune diseases[20–27]. Whether dual TCR T and dual BCR B cells are involved in the immune response to parasitic infections has not been reported. In this study, we leveraged the disruptive advantages of scRNA + TCR + BCR-seq technology in the analysis of dual TCR T and dual BCR B cells to conduct a detailed comparative analysis of the single-cell study data on EG-infected mice by Wu J et al. We innovatively found that dual TCR T and dual BCR B cells may be involved in the immune response and regulation of EG-infected mice, providing a novel perspective for the adaptive immune response mechanisms and new therapeutic targets in parasitic infections.

By comparing the characteristics of dual TCR T and dual BCR B cells in detail, we found that the proportion of dual TCR T cells in the peripheral blood of EG-infected mice was significantly higher than that in uninfected mice, while no significant difference was observed in the spleen. This suggests that dual TCR T cells in peripheral blood may be more involved in the specific response to EG infection. In contrast, the proportion of dual BCR B cells in the peripheral blood of uninfected mice was higher than that in the EG-infected group, suggesting that total single BCR B cells may play a dominant role in the specific response to EG infection in peripheral blood.

The clonal expansion of dual TCR T and dual BCR B cells in the peripheral blood of EG-infected mice was higher than that in uninfected mice, indicating that the clonally expanded dual TCR T and dual BCR B cells in peripheral blood may directly participate in the specific immune response to EG infection. In the spleen, the clonal expansion of dual BCR B cells in the EG-infected group was higher than that in the uninfected group. However, the clonal expansion of dual TCR T cells in the EG-infected group was lower than that in the uninfected group. This may be due to Treg cells in the infected group preventing excessive immune activation. Further comparison of dual TCR Treg cells between the EG-infected and uninfected groups revealed that the expression of inhibitory factors such as PDCD1, Lag3, and CTLA4 was significantly higher in the EG-infected group (Supplementary Figure 3-A)[29–31].

Significant biases in the usage of TRAV and TRBV subfamilies were observed between dual TCR T cells in EG-infected and uninfected mice, as well as biases in the usage of IGHV subfamilies in dual BCR B cells. This suggests that dual TCR T and dual BCR B cells may target different antigenic epitopes of EG in their responses. Additionally, differences in V gene usage were observed between the spleen and peripheral blood in EG-infected mice, indicating tissue-specificity in the antigen recognition sequences of dual TCR T cells during infection (Supplementary Figures 1-C, 1-D,1-E).

Dual TCR T cells in the peripheral blood of EG-infected mice originated from multiple T cell subgroups, including Treg, Th1, Th2, naïve_CD4T, naïve_CD8T, IFN_CD4T, and effector_CD8T. Dual TCR effector_CD8+ T cells exhibited significant clonal expansion (clonetype ≥2). In the spleen of EG-infected mice, dual TCR T cells originated from subgroups excluding Th1 cells, with dual TCR effector_CD8+ T cells, dual TCR Treg cells, and dual TCR IFN_CD4+ T cells showing higher levels of clonal expansion. In contrast, in uninfected mice, clonal expansion was primarily observed in dual TCR naïve CD4 T cells. This suggests that dual TCR effector CD8+ T cells are involved in the specific response to EG infection. Comparative analysis of characteristic gene and cytokine expression between single TCR T and dual TCR T cell subgroups revealed generally consistent expression in infected mice (Supplementary Figures 2-A, 3-A), indicating that single and dual TCR T cells may exert similar immune effects. Treg cells, which express inhibitory transcription factors such as Foxp3, play important roles in immune responses to tumors, autoimmune diseases, and parasitic infections[32–35]. Dual TCR Treg cells have been widely studied [36, 37]. In this study, we found that the proportion of dual TCR Treg cells was higher than that of single TCR Treg cells in both peripheral blood and spleen (Figures 3-B, 4-B). Further analysis revealed that dual TCR Treg cells in the spleen of EG-infected mice exhibited higher expression of inhibitory genes such as Foxp3 and Ikzf2 compared to single TCR Treg cells. This suggests that spleen dual TCR Treg cells may play a primary role in immune regulation, which could be a major direction for future research on parasitic infections. We previously identified potentially effective dual TCR Treg cells in patients with ankylosing spondylitis and Kawasaki disease[38, 39]. These suggest that dual TCR Treg cells may be widely involved in the regulation of immune responses in autoimmune diseases, viral infections, and parasitic infections.

In the early stages of Echinococcus granulosus infection, the immune response is primarily dominated by Th1 cells, which regulate the immune response through transcription factors such as Tbx21 and Stat4, leading to the production of specific cytokines like IFN-γ[40, 41]. In the peripheral blood of EG-infected mice, we observed higher expression of Tbx21 and Stat4 in single TCR Th1 cells compared to dual TCR Th1 cell(s Supplementary Figures 2-A), suggesting that single TCR Th1 cells may play a primary role in the immune response. As EG infection progresses, the immune response shifts to a Th2-dominated response, which primarily involves cells and humoral immunity[42]. Th2 cells mainly secrete IL-4, which promotes Th2-type immune responses, inhibits Th1-type immune responses, and regulates the balance between cell-mediated and humoral immune responses, particularly in combating multicellular parasitic infections such as helminths[43–45]. For example, IL-4 stimulates B cell proliferation and differentiation into antibody-producing cells, particularly increasing the production of IgE and IgG1[46]. In the peripheral blood and spleen of EG-infected mice, we found that single TCR Th2 cells exhibited significantly higher expression of IL-4 compared to dual TCR Th2 cells, suggesting that single TCR Th2 cells play a primary role in immune regulation. High expression of Gata3 in Th2 cells inhibits the production of Th1 cells[47]. In the peripheral blood of EG-infected mice, we observed low expression of Gata3 in dual TCR Th2 cells, while Th1 cell clusters were identified. This suggests that the inhibitory effect of dual TCR Th2 cells may be limited. In contrast, in the spleen of EG-infected mice, dual TCR Th2 cells exhibited high expression of Gata3, and no Th1 cell clusters were identified. This suggests that dual TCR Th2 cells in the spleen may influence Th1 cell differentiation through high expression of Gata3.

Dual BCR B cells in the peripheral blood of EG-infected mice originated from multiple B cell subgroups, including Breg, naïve_B, memory_B, and plasma cells. Dual BCR memory_B cells exhibited higher clonal expansion than dual BCR naïve_B cells. In the spleen of EG-infected mice, dual BCR B cells also included dividing B cells (which were not clustered in peripheral blood), and dual BCR dividing B cells exhibited higher clonal expansion than dual BCR naïve_B cells. In contrast, in uninfected mice, clonal expansion was primarily observed in dual BCR naïve B cells. The high clonal expansion of dual BCR memory_B and dual BCR dividing B cells suggests their potential involvement in the humoral immune response to EG infection. Comparative analysis of clonal expansion in B cell subgroups revealed that single BCR plasma cells exhibited the highest levels of clonal expansion in both spleen and peripheral blood samples. This suggests that single BCR plasma cells are the primary B cell subgroup involved in the immune response to parasitic infections (The role of regulatory B cells in Echinococcus granulosus-infected mice). In the study of gene expression in infected mouse models, it was observed that single BCR B and dual BCR B cells exhibit largely overlapping expression profiles of characteristic genes (Supplementary Figures 2-A , 3-A). This finding suggests that both single BCR B and dual BCR B cells may contribute equivalently to immune responses.

Breg cells, which possess immunoregulatory functions, are involved in immune responses in various diseases, including autoimmune diseases, infections, and tumors. They can suppress inflammatory responses by secreting anti-inflammatory cytokines such as IL-10 and TGF-β[48]. In the peripheral blood and spleen of EG-infected mice, we found that the proportion of dual BCR Breg cells was higher than that of single BCR Breg cells (Figures 3-C, 4-C). Additionally, the proportion of dual BCR Breg cells in the spleen was higher than that in peripheral blood. Further analysis revealed that dual BCR Breg cells in peripheral blood exhibited higher expression of Cd1d1 compared to single BCR Breg cells (Supplementary Figures 2-A). This suggests that dual BCR Breg cells may exert a stronger immunosuppressive effect in the peripheral blood immune environment of EG-infected mice, which could be an important direction for future research on parasitic infections.

KEGG enrichment analysis of single BCR B and dual BCR B cells in the peripheral blood and spleen of EG-infected mice revealed that both single and dual BCR B cells in peripheral blood significantly expressed genes related to “Th1 and Th2 differentiation” and “Antigen processing and presentation.” In the spleen, single and dual BCR B cells significantly expressed genes related to “Intestinal immune network for IgA production” and “Antigen processing and presentation.” This suggests that dual BCR B cells may play an important regulatory role in the specific immune responses of T and B cells during EG infection (Supplementary Figures 7, 8).

Significant overlap in CDR3 sequences was observed between single TCR T and dual TCR T cells, as well as between single BCR B and dual BCR B cells in the peripheral blood and spleen of EG-infected mice. In contrast, minimal overlap was observed in the uninfected group. This suggests that dual TCR T/dual BCR B cells and single TCR T/single BCR B cells may target the same antigenic epitopes and jointly participate in the specific immune response to EG infection. This finding is consistent with our previous observations in autoimmune diseases (ankylosing spondylitis), infections (Kawasaki disease), and tumors (lung cancer), where shared CDR3 sequences were identified between single TCR T and dual TCR T cells[38, 39, 49].

Comparative analysis of the top 10 mRNA molecules expressed in each subgroup of single TCR T and dual TCR T cells, as well as single BCR B and dual BCR B cells, revealed generally consistent expression (Figures 3-D, 4-D). This suggests that dual receptor T or B cells may exert similar immune effects to single receptor T or B cells. However, differences in the expression of some cytokines, cytokine receptors, and transcription factors were observed between the two, suggesting potential differences in the strength of their immune responses and regulation.

By comparing the scRNA + TCR + BCR-seq data of spleen and peripheral blood from EG-infected and uninfected mice in detail, we innovatively found that dual TCR T cells and dual BCR B cells in EG-infected mice may be involved in the specific immune response and regulation to EG infection. Particularly, the clonally expanded dual TCR Treg, dual TCR effector CD8 T, dual BCR Breg, and dual BCR memory/dividing B cells may provide a novel research direction for the complex T and B cell responses in parasitic infections.

## Declaration

### Author contributions

Xinsheng Yao: Conceptualization and design; Kai Quan: Data collection and analysis, drafted the initial version of the manuscript; Peng Su: Participated in date analysis and revision of the manuscript; Hongxia Yang and Yadong Gong: Participated in date analysis. Kai Quan and Peng Su are co-first authors of this article. All authors approved the final version of the manuscript and agree to be accountable for all aspects of the work.

### Funding

This study was supported by the National Natural Science Foundation of China (82160279) and the Guizhou Provincial Hundred level Talent Fund [No. (2018)5637].

### Data availability

The data used in this study are all sourced from public databases (Gene Expression Omnibus repository).scRNA-seq, scTCR-seq, and scBCR-seq data can be found in the Gene Expression Omnibus repository, accession numbers GSE216347 and GSE225312.The data in this database are publicly available under open-access terms, and their use does not require authorization or approval from any institution or regulatory body.The datasets generated and/or analyzed during this study are available from the corresponding author on reasonable request.

### Ethics approval and consent to participate

Not applicable.

### Consent for publication

Not applicable.

### Competing interests

The authors declare no competing interests.

## Reference

1. Zhang W, McManus DP (2008) Vaccination of dogs against Echinococcus granulosus: a means to control hydatid disease? Trends Parasitol 24:419–424. 10.1016/j.pt.2008.05.008

2. Agudelo Higuita NI, Brunetti E, McCloskey C (2016) Cystic Echinococcosis. J Clin Microbiol 54:518–523. 10.1128/JCM.02420-15

3. Armiñanzas C, Gutiérrez-Cuadra M, Fariñas MC (2015) [Hydatidosis: epidemiological, clinical, diagnostic and therapeutic aspects]. Rev Esp Quimioter 28:116–124

4. Sako Y, Nakao M, Nakaya K, et al (2006) Recombinant antigens for serodiagnosis of cysticercosis and echinococcosis. Parasitol Int 55 Suppl:S69–73. 10.1016/j.parint.2005.11.011

5. Albalawi AE, Shater AF, Alanazi AD, et al (2024) High potency of linalool-zinc oxide nanocomposite as a new agent for cystic echinococcosis treatment. Antimicrob Agents Chemother 68:e0173423. 10.1128/aac.01734-23

6. Liu LS, Guo WP, Wang YF, et al (2021) [Hepatic echinococcus granulosus: a clinicopathological analysis of thirteen cases]. Zhonghua Bing Li Xue Za Zhi 50:650–654. 10.3760/cma.j.cn112151-20210202-00119

7. Wang J, Li X, Wang L, et al (2016) A novel long intergenic noncoding RNA indispensable for the cleavage of mouse two-cell embryos. EMBO Rep 17:1452–1470. 10.15252/embr.201642051

8. Vuitton DA (2004) Echinococcosis and allergy. Clin Rev Allergy Immunol 26:93–104. 10.1007/s12016-004-0004-2

9. McNab F, Mayer-Barber K, Sher A, et al (2015) Type I interferons in infectious disease. Nat Rev Immunol 15:87–103. 10.1038/nri3787

10. Harraga S, Godot V, Bresson-Hadni S, et al (1999) Clinical efficacy of and switch from T helper 2 to T helper 1 cytokine profile after interferon alpha2a monotherapy for human echinococcosis. Clin Infect Dis 29:205–206. 10.1086/520157

11. Zhang S, Zhou Y, Su L, et al (2017) In vivo evaluation of the efficacy of combined albedazole-IFN-α treatment for cystic echinococcosis in mice. Parasitol Res 116:735–742. 10.1007/s00436-016-5339-0

12. Wang J, Müller S, Lin R, et al (2017) Depletion of FoxP3+ Tregs improves control of larval Echinococcus multilocularis infection by promoting co-stimulation and Th1/17 immunity. Immun Inflamm Dis 5:435–447. 10.1002/iid3.181

13. Qi X, Shan J, Liu X, et al (2021) The role of regulatory B cells in Echinococcus granulosus-infected mice. Parasitol Res 120:1389–1404. 10.1007/s00436-020-07025-3

14. Padovan E, Casorati G, Dellabona P, et al (1993) Expression of two T cell receptor alpha chains: dual receptor T cells. Science 262:422–424. 10.1126/science.8211163

15. Vettermann C, Schlissel MS (2010) Allelic exclusion of immunoglobulin genes: models and mechanisms. Immunol Rev 237:22–42. 10.1111/j.1600-065X.2010.00935.x

16. Outters P, Jaeger S, Zaarour N, Ferrier P (2015) Long-Range Control of V(D)J Recombination & Allelic Exclusion: Modeling Views. Adv Immunol 128:363–413. 10.1016/bs.ai.2015.08.002

17. Triebel F, Breathnach R, Graziani M, et al (1988) Evidence for expression of two distinct T cell receptor beta-chain transcripts in a human diphtheria toxoid-specific T cell clone. J Immunol 140:300–304

18. Malissen M, Trucy J, Letourneur F, et al (1988) A T cell clone expresses two T cell receptor alpha genes but uses one alpha beta heterodimer for allorecognition and self MHC-restricted antigen recognition. Cell 55:49–59. 10.1016/0092-8674(88)90008-6

19. Rezanka LJ, Kenny JJ, Longo DL (2005) 2 BCR or NOT 2 BCR - receptor dilution: a unique mechanism for preventing the development of holes in the protective B cell repertoire. Immunobiology 210:769–774. 10.1016/j.imbio.2005.10.008

20. Sarukhan A, Garcia C, Lanoue A, von Boehmer H (1998) Allelic inclusion of T cell receptor alpha genes poses an autoimmune hazard due to low-level expression of autospecific receptors. Immunity 8:563–570. 10.1016/s1074-7613(00)80561-0

21. Padovan E, Casorati G, Dellabona P, et al (1995) Dual receptor T-cells. Implications for alloreactivity and autoimmunity. Ann N Y Acad Sci 756:66–70. 10.1111/j.1749-6632.1995.tb44482.x

22. Schuldt NJ, Binstadt BA (2019) Dual TCR T Cells: Identity Crisis or Multitaskers? J Immunol 202:637–644. 10.4049/jimmunol.1800904

23. Fournier EM, Velez M-G, Leahy K, et al (2012) Dual-reactive B cells are autoreactive and highly enriched in the plasmablast and memory B cell subsets of autoimmune mice. J Exp Med 209:1797–1812. 10.1084/jem.20120332

24. Corthay A, Nandakumar KS, Holmdahl R (2001) Evaluation of the percentage of peripheral T cells with two different T cell receptor alpha-chains and of their potential role in autoimmunity. J Autoimmun 16:423–429. 10.1006/jaut.2001.0504

25. Sang A, Danhorn T, Peterson JN, et al (2018) Innate and adaptive signals enhance differentiation and expansion of dual-antibody autoreactive B cells in lupus. Nat Commun 9:3973. 10.1038/s41467-018-06293-z

26. Vincent BG, Serody JS (2013) One is better than two: TCR pairing and GVHD. Sci Transl Med 5:188fs21. 10.1126/scitranslmed.3006431

27. Ji Q, Perchellet A, Goverman JM (2010) Viral infection triggers central nervous system autoimmunity via activation of CD8+ T cells expressing dual TCRs. Nat Immunol 11:628–634. 10.1038/ni.1888

28. The Clonal Selection Theory of Acquired Immunity. | JAMA Internal Medicine | JAMA Network. https://jamanetwork.com/journals/jamainternalmedicine/article-abstract/564625. Accessed 10 Apr 2025

29. Kumar P, Bhattacharya P, Prabhakar BS (2018) A comprehensive review on the role of co-signaling receptors and Treg homeostasis in autoimmunity and tumor immunity. J Autoimmun 95:77–99. 10.1016/j.jaut.2018.08.007

30. Kim MJ, Kim K, Park HJ, et al (2023) Deletion of PD-1 destabilizes the lineage identity and metabolic fitness of tumor-infiltrating regulatory T cells. Nat Immunol 24:148–161. 10.1038/s41590-022-01373-1

31. Huang C-T, Workman CJ, Flies D, et al (2004) Role of LAG-3 in regulatory T cells. Immunity 21:503–513. 10.1016/j.immuni.2004.08.010

32. Wu X, Zhou Z, Cao Q, et al (2023) Reprogramming of Treg cells in the inflammatory microenvironment during immunotherapy: a literature review. Front Immunol 14:1268188. 10.3389/fimmu.2023.1268188

33. Togashi Y, Shitara K, Nishikawa H (2019) Regulatory T cells in cancer immunosuppression - implications for anticancer therapy. Nat Rev Clin Oncol 16:356–371. 10.1038/s41571-019-0175-7

34. Mikami N, Kawakami R, Sakaguchi S (2020) New Treg cell-based therapies of autoimmune diseases: towards antigen-specific immune suppression. Curr Opin Immunol 67:36–41. 10.1016/j.coi.2020.07.004

35. Zhang Q, Ye J, Zheng H (2016) Dexamethasone attenuates echinococcosis-induced allergic reactions via regulatory T cells in mice. BMC Immunol 17:4. 10.1186/s12865-016-0141-4

36. Miao J, Zhu P (2018) Functional Defects of Treg Cells: New Targets in Rheumatic Diseases, Including Ankylosing Spondylitis. Curr Rheumatol Rep 20:30. 10.1007/s11926-018-0729-1

37. Tuovinen H, Salminen JT, Arstila TP (2006) Most human thymic and peripheral-blood CD4+ CD25+0020regulatory T cells express 2 T-cell receptors. Blood 108:4063–4070. 10.1182/blood-2006-04-016105

38. Yuanyuanxu null, Qipeng null, Qingqingma null, Yao X (2024) scRNA + TCR-seq revealed the dual TCR pTh17 and Treg T cells involvement in autoimmune response in ankylosing spondylitis. Int Immunopharmacol 135:112279. 10.1016/j.intimp.2024.112279

39. Xu Y, Yuan Y, Mou L, et al (2024) scRNA+TCR-seq reveals the pivotal role of dual receptor T lymphocytes in the pathogenesis of Kawasaki disease and during IVIG treatment. Front Immunol 15:1457687. 10.3389/fimmu.2024.1457687

40. Butcher MJ, Zhu J (2021) Recent advances in understanding the Th1/Th2 effector choice. Fac Rev 10:30. 10.12703/r/10-30

41. Zhang Y, Zhang Y, Gu W, Sun B (2014) TH1/TH2 cell differentiation and molecular signals. Adv Exp Med Biol 841:15–44. 10.1007/978-94-017-9487-9_2

42. Early peritoneal immune response during Echinococcus granulosus establishment displays a biphasic behavior - PubMed. https://pubmed.ncbi.nlm.nih.gov/21912714/. Accessed 10 Apr 2025

43. Wang C, Yang S-H, Niu N, et al (2021) lncRNA028466 regulates Th1/Th2 cytokine expression and associates with Echinococcus granulosus antigen P29 immunity. Parasit Vectors 14:295. 10.1186/s13071-021-04795-2

44. Kubo M (2021) The role of IL-4 derived from follicular helper T (TFH) cells and type 2 helper T (TH2) cells. Int Immunol 33:717–722. 10.1093/intimm/dxab080

45. Golubovskaya V, Wu L (2016) Different Subsets of T Cells, Memory, Effector Functions, and CAR-T Immunotherapy. Cancers (Basel) 8:36. 10.3390/cancers8030036

46. Kopf M, Le Gros G, Bachmann M, et al (1993) Disruption of the murine IL-4 gene blocks Th2 cytokine responses. Nature 362:245–248. 10.1038/362245a0

47. Zhu J, Yamane H, Cote-Sierra J, et al (2006) GATA-3 promotes Th2 responses through three different mechanisms: induction of Th2 cytokine production, selective growth of Th2 cells and inhibition of Th1 cell-specific factors. Cell Res 16:3–10. 10.1038/sj.cr.7310002

48. Catalán D, Mansilla MA, Ferrier A, et al (2021) Immunosuppressive Mechanisms of Regulatory B Cells. Front Immunol 12:611795. 10.3389/fimmu.2021.611795

49. Peng Q, Xu Y, Yao X (2024) scRNA+ TCR-seq revealed dual TCR T cells antitumor response in the TME of NSCLC. J Immunother Cancer 12:e009376. 10.1136/jitc-2024-009376

